# Structure of BrlR reveals a potential pyocyanin binding site

**DOI:** 10.1101/188136

**Authors:** Harikiran Raju, Rukmini Sundararajan, Rohan Sharma

## Abstract

The transcriptional regulator BrlR from Pseudomonas aeruginosa is a member of the MerR family of multidrug transport activators. Studies have shown BrlR plays an important role in high level drug tolerance of P. aeruginosa in biofilm. Its drug tolerance ability can be enhanced by 3′,5′-cyclic diguanylic acid (c-di-GMP). Here, we show the apo structure of BrlR and the direct binding between GyrI-like domain of BrlR and P. aeruginosa toxin pyocyanin. Furthermore, pyocyanin can enhance the binding between BrlR and DNA in vitro. These findings suggest BrlR can serve as the binding partner for both c-di-GMP and pyocyanin.

## Introduction

*P. aeruginosa* infections have complex pathophysiology and are difficult to eliminate [1]. Its intrinsic ability to develop resistance to antibiotics, formation of impenetrable biofilms and release of a large quantity of virulence factors all contribute to *P. aeruginosa* infections. In addition to secreted attacking protein enzymes, including elastase, alkaline protease and LasA protease, many secondary metabolites also are released, which can cause serious damage to the host.

Pyocyanin is a blue redox-active secondary metabolite, which can easily penetrate biological membranes and cause damages [2, 3]. It is synthesized from chorismate through a series of complex steps mediated by gene products encoded by two phzABCDEFG operons and its precursors will be modified into the tricyclic compound by the phzH, phzM and phzS genes [4, 5]. The synthesis of pyocyanin is regulated by quorum sensing (QS) systems, but the details are not clear.

Pyocyanin was reported to kill fungi and Caenorhabditis elegans [6, 7]. Recent studies demonstrated that pyocyanin is required for lung infection in mice [8, 9]. Moreover, large quantities of pyocyanin can be readily recovered from the sputum of patients with CF infected by *P. aeruginosa* [10]. It has been reported that pyocyanin interferes with multiple cellular functions that it can cause oxidative stress, inactivate of V-ATPase, decrease mitochondrial and cytoplasmic aconitase activity and ATP levels [2, 11]. Many researches about pyocyanin focus on its role in pathogenesis, yet the regulation and signal transduction pathway it participates have long been ignored. Until now, only few pyocyanin receptors are reported. The arylhydrocarbonreceptor (AhR) from human is a highly conserved ligand-dependent transcription factor, which can sense pyocyanin, induce detoxifying enzymes and modulate immune cell differentiation and responses [12-14]. RmcA from *P. aeruginosa* is a multiple domain protein with both phosphodiesterase (PDE) and diguanylate cyclase (DGC) domain, whose activity is modulated by pyocyanin through its PAS domain [15].

BrlR protein is a transcription regulator in *P. aeruginosa.* It can only be detected in biofilm cells and activated by bacterial secondary messenger c-di-GMP [16]. The activated BrlR can upregulate the expression of multidrug efflux pumps, such as MexAB-OprM and MexEF-OprN efflux pumps. The knock-out of BrlR leads to the susceptibility of the biofilm when the bacteria were treated with five different classes of antibiotics [17]. BrlR belongs to the MerR protein family and consists of three domains with the middle coiled-coil region flanked by N-terminal HTH_MerR domain and GyrI-like domain at the C-terminus. MerR protein family have been demonstrated as multidrug binding (MDR) proteins, and some of them can bind potential multidrug molecules directly. In our previous study, we determined the complex structure of BrlR-c-di-GMP and identified one conserved drug-binding pocket. However, so far, there is no report regarding which small molecule can bind into this pocket and what is the common link between biofilm and multidrug resistance in *P. aeruginosa*. In this study, we demonstrate that BrlR from *P. aeruginosa* is binding partner for pyocyanin and identify the GyrI-like domain as potential binding site for pyocyanin.

## Materials and Methods

### Protein expression and purification

The expression and purification of full length *brlR* gene and 119-end fragment (GyrI-like domain) were the same as previous protocol [18]. Briefly, the genes encoding the corresponding sequences were amplified by polymerase chain reaction (PCR) and inserted into the pET-28a vector (Novagen). The *E. coli* BL21 (DE3) cells were used as host for expression. The proteins were obtained by two-step purification procedure (Ni-affinity and gel filtration purification). The N-terminal His-tag of BrlR removed by thrombin protease before gel filtration was used for crystallization and The N-terminal His-tag of BrlR left intact through the purification was used for western blot (WB). The protein was then concentrated in gel filtration buffer (20 mM Tris pH 7.5, 150 mM NaCl, 5% Glycerol). Protein purity was checked by Coomassie Brilliant Blue-stained sodium dodecyl sulfate polyacrylamide gel electrophoresis (SDS-PAGE). The mutagenesis of BrlR (W150A) was performed by overlap-extension PCR and the expression and purification of these mutant proteins were the same as the wild type.

### Crystallization, data collection, structure determination and refinement

Purified BrlR proteins were concentrated to approximately 10 mg/mL in the gel filtration buffer and screened for crystallization using commercial available kits (Molecular Dimension). Sitting-drop vapor diffusion method was used in 24-well Itelli-plate to optimize the crystals at 20 °C under condition (10% v/v 1,4-Dioxane, 0.1 M MES/NaOH pH 6.5, 1.6 M Ammonium sulfate)( Structure Screen 1–2, Molecular Dimension). The grown crystals were dehydrated by 1.8 M Lithium sulfate to improve the resolution.

The diffraction data were collected at the beamline BM14 at the European Synchrotron Radiation Facility (ESRF). Macromolecular crystal annealing (MCA) was performed. Briefly, before data collection, the cold nitrogen stream was blocked about 3 s, three times, with intervals of 5 s[19]. The data were indexed, integrated and scaled with HKL2000 [20]. Phaser (PHENIX) was used to do the molecular replacement [21]. The coordinate of BrlR from c-di-GMP complex structure (PDB: 5XQL) was used as the search model. Coot was used to modify the result model and phenix.refine was used for refinement [21, 22]. The structure of BrlR was deposited in RCSB with a PDB code 5YC9. Crystallographic statistics are summarized in Table 1. All the structure figures were prepared with PyMOL (http://www.pymol.org).

**Table 1.** Data collection and refinement statistics

|  |  |  |  |
| --- | --- | --- | --- |
| **Data collection** |  |  |  |
| Wavelength (Å) | 0.98 |  |  |
| Space group | P 65 |  |  |
| *a*, *b*, *c* (Å) | 111.64 | 111.64 | 261.13 |
| $\alpha$, $\beta$, $\gamma$ (°) | 90.00 | 90.00 | 120.00 |
| Resolution range (Å) | 35.00-2.90 (2.95-2.90)* |  |  |
| Completeness (%) | 99.7 (100.0) |  |  |
| Redundancy | 7.9 (7.9) |  |  |
| $\langle I/\sigma(I) \rangle$ | 18.71 (1.39) | | |
| Rpim. | 0.037 (0.606) |  |  |
| CC1/2 | 0.564 |  |  |
| Overall B factor from Wilson plot (Å<sup>2</sup>) | 43.16 |  |  |
| **Refinement** |  |  |  |
| No. reflections | 60294 |  |  |
| Final Rcryst /Final Rfree | 0.2693/0.2969 |  |  |
| *No. of non-H atoms* |  |  |  |
| Protein | 8290 |  |  |
| *R.m.s. deviations* |  |  |  |
| Bonds (Å) | 0.006 |  |  |
| Angles (°) | 1.154 |  |  |
| *Average B factors (Å<sup>2</sup>)* |  |  |  |
| Protein | 65.84 |  |  |
| *Ramachandran plot* |  |  |  |
| Most favoured (%) | 91.21 |  |  |
| Allowed (%) | 8.59 |  |  |
| Outliers (%) | 0.20 |  |  |

\*Values in parentheses correspond to highest resolution shell.

### Electrophoretic mobility shift assay (EMSA)

According to previous papers, three DNA probes are designed and labeled with Cyanine 5 (Cy5) fluorescent dye (Thermofisher Scientific) (Supplementary table1) [17, 23]. The DNA probes (0.5 pmol), BrlR proteins (25 pmol) and different concentration of pyocyanin (Everon Life Sciences) or c-di-GMP (Labex Corporation) were incubated for 30 min at 25°C in binding buffer (10 mM Tris-HCl pH 7.5, 50 mM KCl, 1 mM EDTA, 1 mM DTT and 5% glycerol) with a total volume of 20 μl Then, the samples were subjected to electrophoresis with a 6% polyacrylamide glycine gel using 0.5×TBE running buffer on ice for 80 min. Imaging and data analyses were performed on a LI-COR imaging system (LI-COR Biosciences).

### Ligand trapping assay

In order to check the binding between BrlR and pyocyanin, the ligand trapping assay were performed with a 5 KD cutoff centrifugal filter. BrlR proteins were incubated with its ligand in a 1:2 ratio for 20 min at RT. Then, BrlR proteins with ligand were concentrated by centrifugal filter at 4 °C for 30 min. The flow-through was collected to measure the concentration of un-trapped ligand at the wavelength of 310 nm and the buffer with ligand was used as control experiment. Every binding assay was repeated at least three times.

### Binding affinity measurement

The T_m_ value of BrlR was determined by thermal-denaturation assay with 2 °C increment from 50 °C to 68 °C. Briefly, 20 μg protein was incubated at denaturing temperature for 3 min followed by centrifugation for 5 min. Then, the supernatant was collected for concentration measurement. The Tm (57.5 °C) was obtained by fitting the denaturation curve. The BrlR-pyocyanin binding affinity was measured by ligand concentration-dependent thermo-denaturation assay. The 0.05 μM BrlR protein was incubated with various concentrations of pyocyanin (0.5 μM, 1 μM, 5 μM, 10 μM, 20 μM) and denatured at 57.5 °C for 3 min. The supernatants were collected for western blot (WB). The protein was detected by using primary mouse anti-His antibody (1:5000, Clontech) and secondary HRP conjugated Rabbit anti-mouse antibody (1:2000, Cell signaling). The image was analyzed using image J and Kd was calculated with fitted curve.

## Result

### Tetrameric apo BrlR structure

The crystal structure of apo BrlR was determined at 3.3 Å, with four BrlR molecules in the asymmetry unit (AU). The density is poor around two regions (residues 27aa-40aa and 137aa-144aa) in each protomer because these regions are flexible (Fig. 1A). According to previous BrlR-c-di-GMP complex structure, residues (27aa-40aa) take part in c-di-GMP binding and residues (137aa-144aa) are located near the drug-binding pocket [18]. Therefore, both areas show the binding potential for small molecules due to their flexibility.

**Figure 1.**
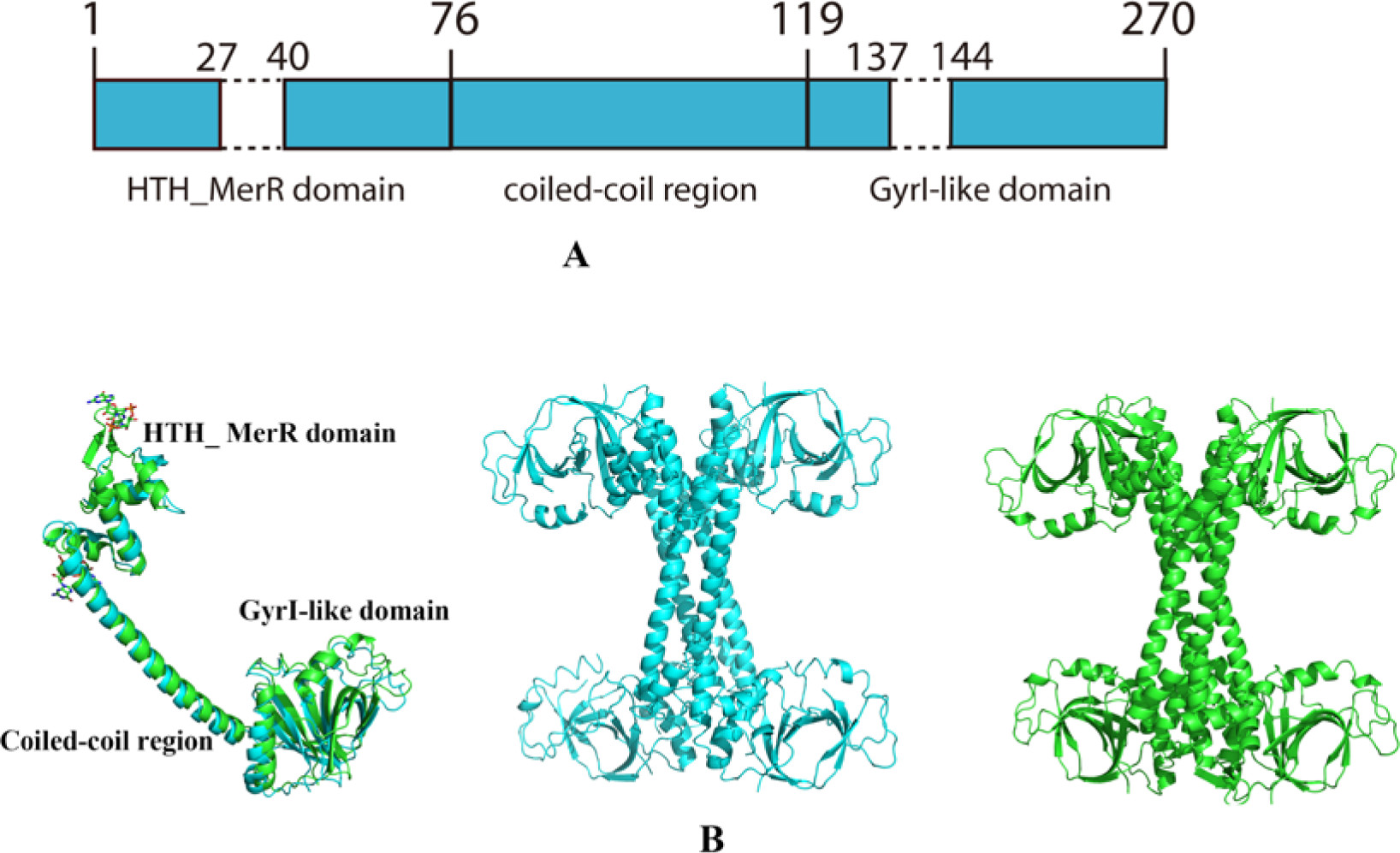
Overall structure of apo BrlR structure. (A) Domain structure of BrlR. Dash line indicates the disordered regions in our apo BrlR structure. (B) Left panel: Superposition of cartoon presentation of apo BrlR monomer with BrlR monomer from c-di-GMP complex (PDB: 5XQL). No obvious conformation change is observed between the two forms. Apo BrlR monomer can be clearly divided into three parts: N-terminal HTH_ MerR domain (1-76 aa), middle coiled-coil region (77-119 aa) and C-terminal GyrI-like domain (120-270 aa). Middle and right panel: cartoon presentation of tetramer of apo BrlR (in cyan) and BrlR tetramer from c-di-GMP bound form (in green). Just like the c-di-GMP bound form, the HTH_ MerR domain, middle coiled-coil region and GyrI-like domain all take part in the massive interaction among the four BrlR protomers in the apo form, while the relative position of these four chains has only a minor difference.

The overall structure of apo BrlR is much similar to the c-di-GMP bound form. No dramatic conformation difference is observed after c-di-GMP binding and some minor differences may be due to the crystal packing artifact (Fig. 1B). Like the c-di-GMP bound form, apo BrlR structure can be clearly divided into three parts: N-terminal HTH_ MerR domain, middle coiled-coil region and C-terminal GyrI-like domain (Fig. 1). The apo BrlR exists as tetramer in solution which have been revealed by previous gel filtration experiment [18].

The c-di-GMP binding did not change the overall conformation and oligomerization state of BrlR (Fig. 1B). Therefore, how the c-di-GMP stimulates the BrlR remains elusive.

### Pyocyanin binds BrlR in the GyrI-like domain

The similarity and conservation of GyrI-like domain between BrlR and other MDR proteins discussed in previous paper indicates BrlR can bind flat shape molecules (Fig. S1). For detailed analysis, we compared the GyrI-like domain of BrlR with SAV2435 (PDB code: 5KAU). The GyrI-like domain of BrlR and SAV2435 align well, with a root-mean-square deviation (RMSD) value of 2.3 Å over 165 residues (Fig. 2A). Like SAV2435, the drug-binding pocket of BrlR GyrI-like domain is deep and hydrophobic, which indicates that this site may have the same preference for small molecules with aromatic rings. Unlike BmrR and SAV2435, which are reported to bind many molecules, no small molecules are reported to bind the GyrI-like domain of BrlR so far. We noticed that the central xanthene ring of rhodamine 6G (RH6G) in SAV2435 complex structure is very similar to pyocyanin, which exists in large quantities in *P. aeruginosa*.

**Figure 2.**
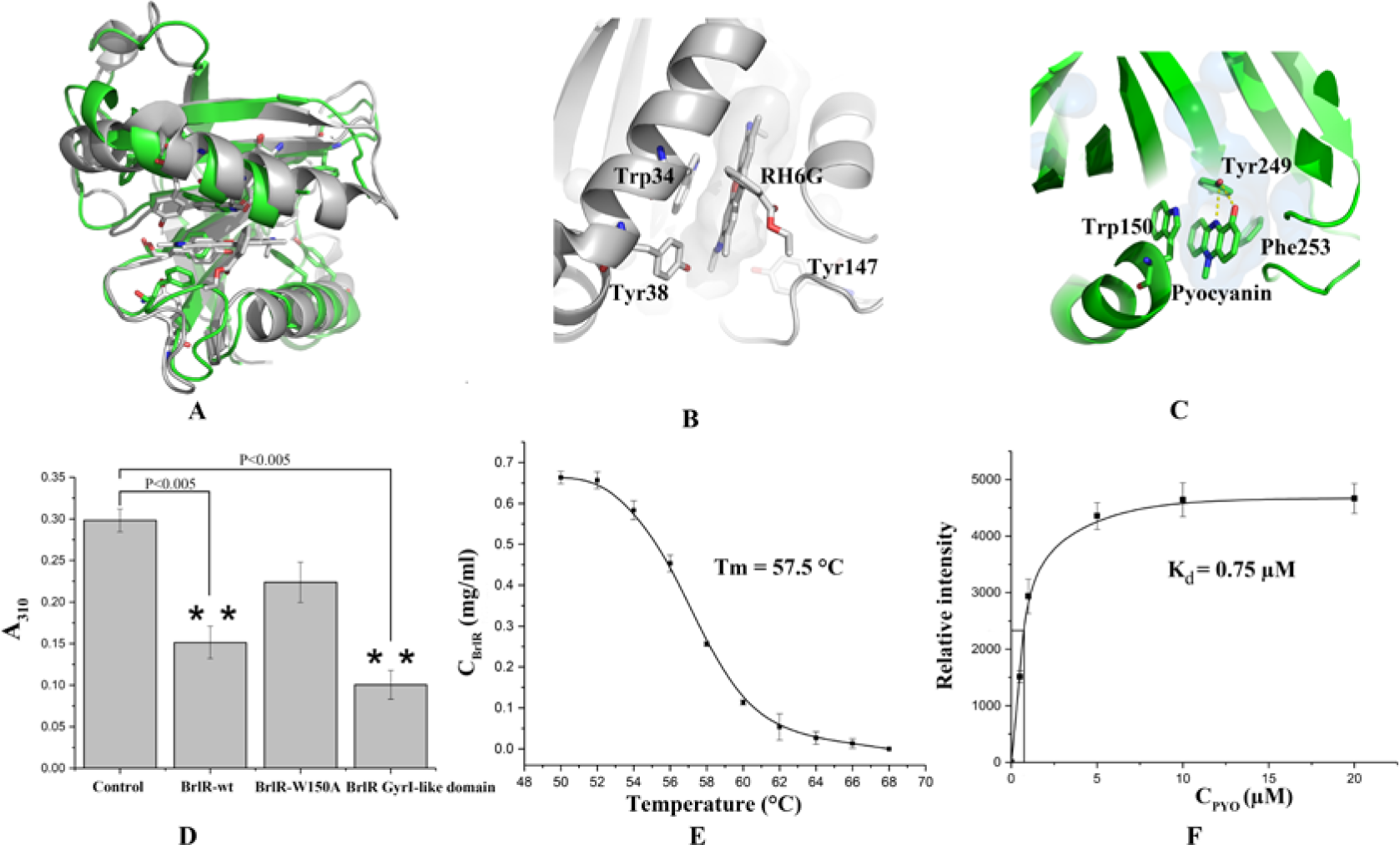
Pyocyanin binds BrlR in the GyrI-like domain. (A) Superposition of cartoon presentation of GyrI-like domain of BrlR with RH6G bound SAV2435 (PDB: 5KAU). BrlR is in cyan and SAV2435 is in grey. Aromatic residues around the drug-binding pocket and RH6G molecule are rendered as stick. (B) RH6G binding site in SAV2435. The binding cavity is presented by surface; key residues and RH6G are rendered as stick. (C) Predicted pyocyanin binding site in BrlR GyrI-like domain. The potential binding cavity is presented by surface; predicted key residues and pyocyanin are rendered as stick. The yellow dash lines indicate the possible interaction between pyocyanin Tyr249, which may play a role in binding specificity. (D) Ligand trapping assay for BrlR and pyocyanin. While the GyrI-like domain of BrlR can even bind pyocyanin tighter, the full-length W150A mutation binds less pyocyanin. Experiments were carried out at least in triplicate. Error bars indicate standard deviation. **Significantly different from control; P < 0.005. P-values are calculated by two-tailed Student’s t-test method. (E) Denaturation curve of BrlR. Error bars indicate standard deviation. (F) Ligand concentration-dependent thermal-denaturation assay for BrlR and pyocyanin. Data are calculated on relative intensity of bands detected by WB and analysis using ImageJ. Kd stands for the dissociate constant. Error bars indicate standard deviation.

It is reported that exogenous added pyocyanin can affect the expression of dozens of genes [24]. It may be due to the chemical property of pyocyanin served as a redox-active molecule or its directly binding to some transcription regulators that cause this regulation. However, no transcription regulators are identified as pyocyanin receptors so far. We noticed that among the upregulated genes, eight of which encode putative transporters (i.e. the RND transporters mexGHI-opmD, PA3923-3922-opmE and the putative Major Facilitator Superfamily transporter PA3718). The BrlR gene expression level is also upregulated with 2.3 fold. We found this regulation pattern is somewhat similar to the BrlR gene does. In addition to itself, BrlR will activate genes encoding the MexAB-OprM and MexEF-OprN multidrug efflux pumps after stimulation by c-di-GMP [17]. In such condition, we hypothesize that BrlR can bind pyocyanin and trigger its gene regulation function.

To better characterize the binding property of BrlR towards pyocyanin, we modelled the pyocyanin into the surface cavity of GyrI-like domain in silicon according to the binding mode of RH6G in SAV2435 (Fig. 2B and C). The result shows both RH6G and pyocyanin are buried in the cavity of the drug-binding pocket, with several aromatic residues around them. Like Trp34 in SAV2435, Trp150 form stack interaction with pyocyanin in this model. At the same time, Tyr249 form hydrogen bond with pyocyanin, which will play a role in its binding specificity (Fig. 2C). In order to confirm this hypothesis, we tested the binding between BrlR protein and pyocyanin using ligand trapping assay. As is shown in Fig. 2D, it clearly show that the BrlR can bind the pyocyanin and reduce its chance to flow through the filter membrane. The binding also change solution color from blue to green especially in the more concentrated condition (Fig. S2). The Tm value of BrlR is about 57.5 °C and 20 μM pyocyanin can increase it to 61.5 °C (Fig. 2E and S2C). To test whether pyocyanin binding pocket is located in our predicted site in the C-terminal GyrI-like domain, we used the C-terminal part of BrlR for this binding assay and it shows it binds pyocyanin much better than full length protein, which indicates the full length protein may be in a more regulated conformation (Fig. 3D). The binding affinity between BrlR and pyocyanin is about 0.75 μM (Fig. 2F and S2D), which is comparable to the c-di-GMP binding affinity (2.2 μM) [16]. However, the binding between W150A mutation protein and pyocyanin is decreased as is shown by ligand trapping assay (Fig. 2D), which suggests the predicted binding site is correct.

**Figure 3.**
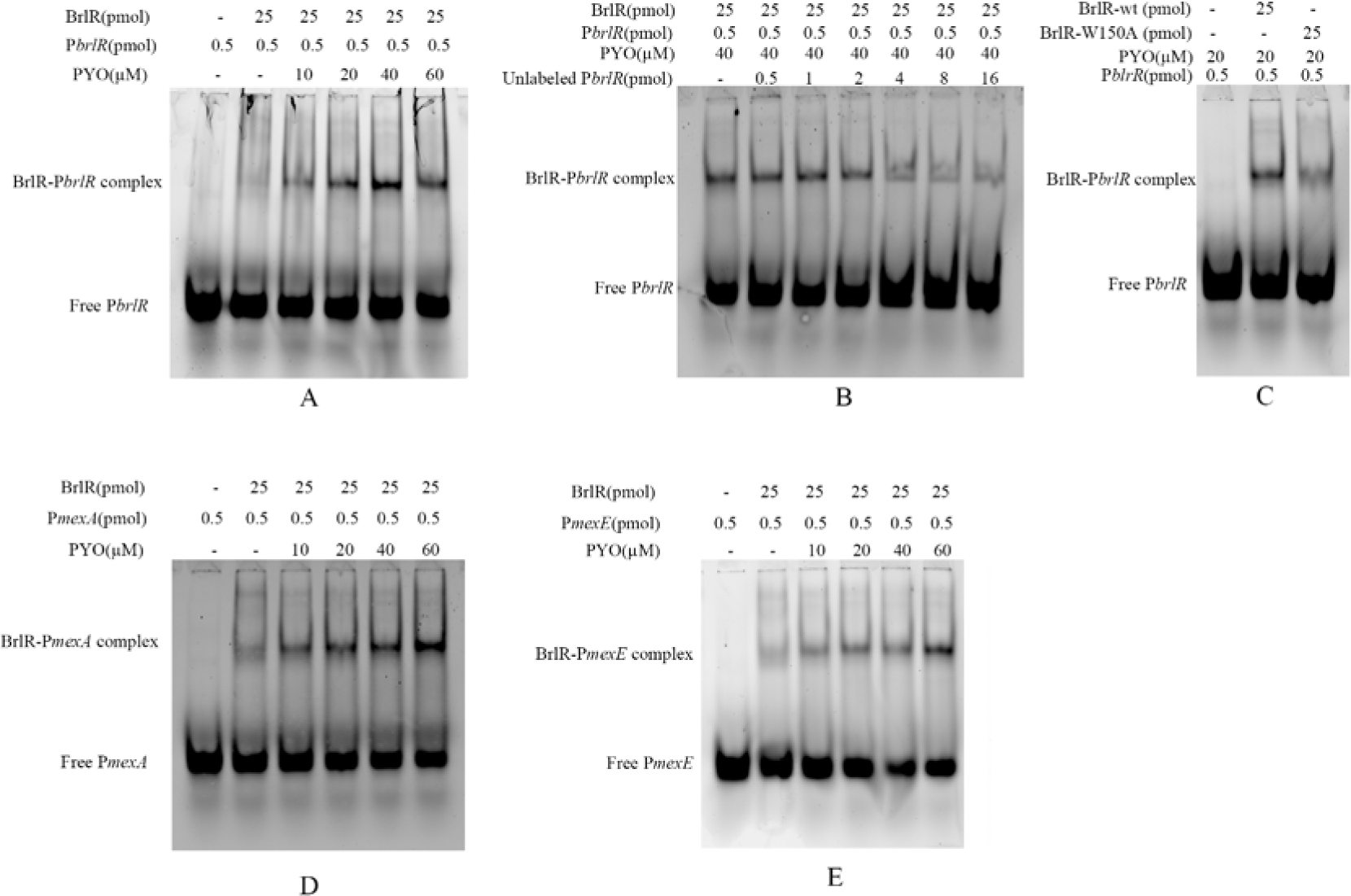
EMSA assays for BrlR with pyocyanin. (A) EMSA assays with P*brlR*. The basic DNA binding of apo BrlR is relatively weak and BrlR-P*brlR* complex can be clearly observed only after adding 10 μM pyocyanin. Our assays shows that increased pyocyanin concentration can enhance the DNA binding of BrlR. (B) EMSA assays with unlabeled P*brlR*. With the increase of unlabeled P*brlR*, the bound labeled P*brlR* continues to decrease. (C) EMSA assays for BrlR W150A mutation. The binding between W150A mutation and P*brlr* promoter can not be enhanced by pyocyanin. (D, E) EMSA assays with P*mexA* and P*mexE*. The effect of pyocyanin on BrlR binding towards these two fragments are similar with P*brlR*. (F) EMSA assays for BrlR with pyocyanin and c-di-GMP. pyocyanin and c-di-GMP can cooperatively enhance the binding ability of BrlR to DNA. All of the assays were performed at least three times.

### Pyocyanin can enhance the DNA binding of BrlR

It was reported that BrlR can bind to its own promoter (P*brlR*) and the promoter of *mexA* (P*mexA*) and *mexE* (P*mexE*) [17]. While c-di-GMP can enhance this binding, we speculate that pyocyanin could function in the same way. To characterize the effect of pyocyanin binding, we performed the EMSA assays of BrlR with Cy5 labeled P*brlR*, P*mexA* and P*mexE* under different concentration of pyocyanin (Fig. 3). Due to the chemical activity, we checked the effect of pyocyanin on Cy5 labeled DNA (Fig. S3). It seems that the pyocyanin can quench the Cy5 fluorescent signals especially at high concentration. While the DNA binding of apo BrlR is weak, it is clearly shown that increased pyocyanin concentration can enhance the binding of BrlR towards P*brlR*, P*mexA* and P*mexE*. Because BrlR can non-specifically bind many dyes (data not shown), we use unlabeled PbrlR, PmexA and PmexE to show the specificity (Fig. 3B and S4A and C). In our assays, high concentration of unlabeled PbrlR, PmexA and PmexE can successfully compete the BrlR protein, which indicates the labelling dye does not disturb our EMSA assays. The failure of obtaining the complex structure of BrlR and pyocyanin makes the activation mechanism still unknown. In our size-exclusion chromatography (SEC) experiments, the apo and pyocyanin bound form BrlR share the same elution time, which suggests that pyocyanin does not change the oligomeric sate of BrlR, just like the c-di-GMP (Fig. S5). The W150A mutation which lost the pyocyanin binding shows a compromised binding ability to PbrlR, PmexA and PmexE DNA in the presence of pyocyanin (Fig. 3C and S4B and D). These data shown above indicate the pyocyanin can bind and regulate the transcription factor BrlR.

## Discussion

The secondary messenger c-di-GMP can promote the biofilm formation in *P. aeruginosa* and the BrlR protein can only be detected after the biofilm formation. The binding of pyocyanin or c-di-GMP by BrlR leads to the expression of *BrlR* gene and BrlR regulated multidrug efflux pump genes [17]. It seems that both these two small molecules contribute to the multidrug resistance of *P. aeruginosa*. Due to the lacking of BrlR-ligand-DNA ternary complex structure, the molecular mechanism of BrlR stimulation by c-di-GMP or pyocyanin remains elusive.

In summary, our data suggest that BrlR is the binding partner for both c-di-GMP and pyocyanin. As an unusual transcription regulator, BrlR has involved two separate binding site towards c-di-GMP and pyocyanin. The next step study will focus on the physiological significance of this binding between BrlR and pyocyanin. It also shows the BrlR will be a promising drug target for the cure of intractable infection of *P. aeruginosa*.

## Acknowledgements

The authors acknowledge the staff of BM14 beamline at the European Synchrotron Rediation Facility (ESRF) for their support in the data collection. The authors acknowledge financial support from University of Calcutta.

